## Supplemental Materials for "The Goldilocks Effect: Female geladas in mid-sized groups have higher fitness"

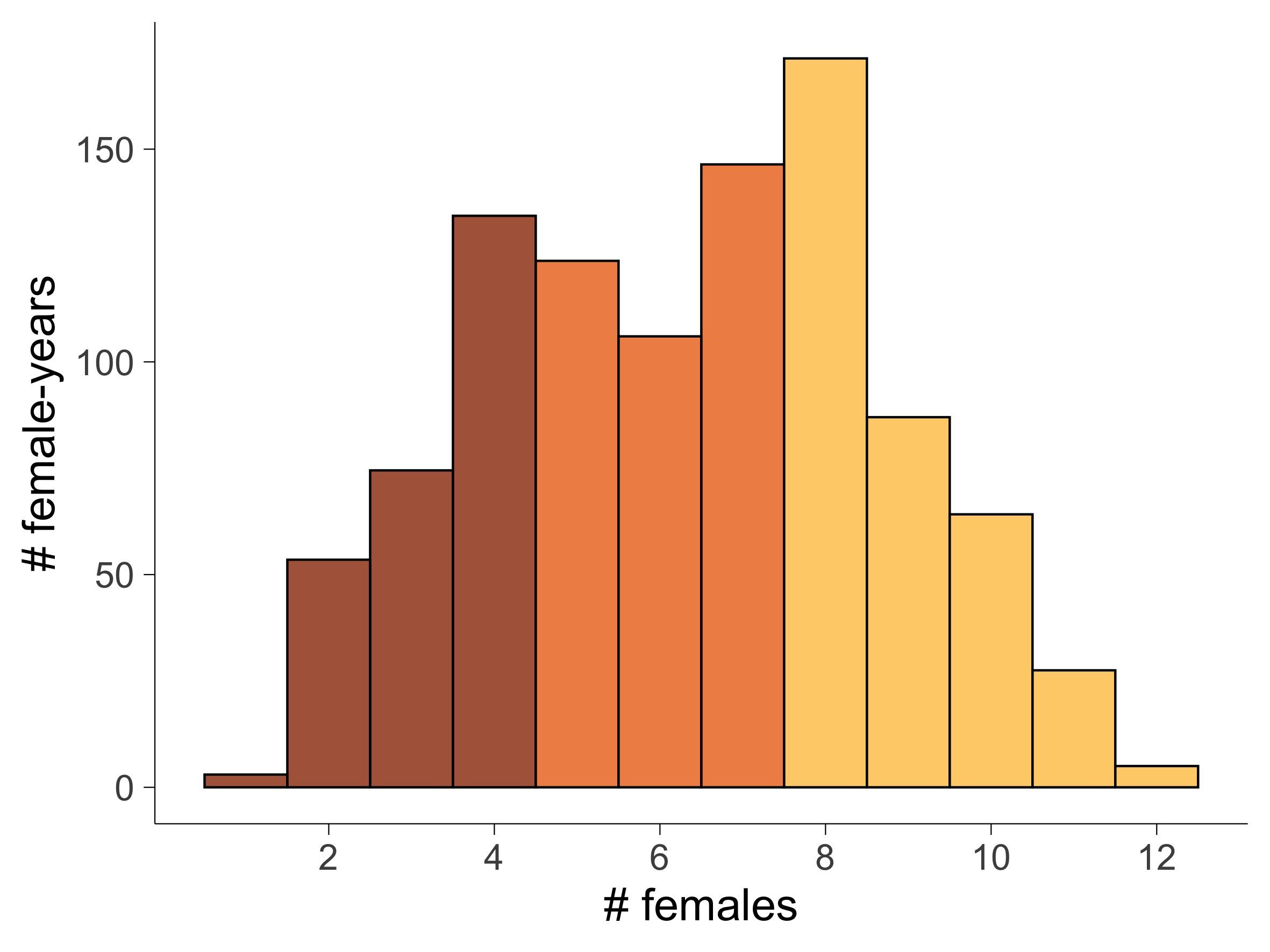

**Figure S1. Distribution of unit sizes across the dataset.** The total number of female-years at each unit size in the dataset. Red, orange, and yellow bins denote small, mid-sized, and large unit size categories.

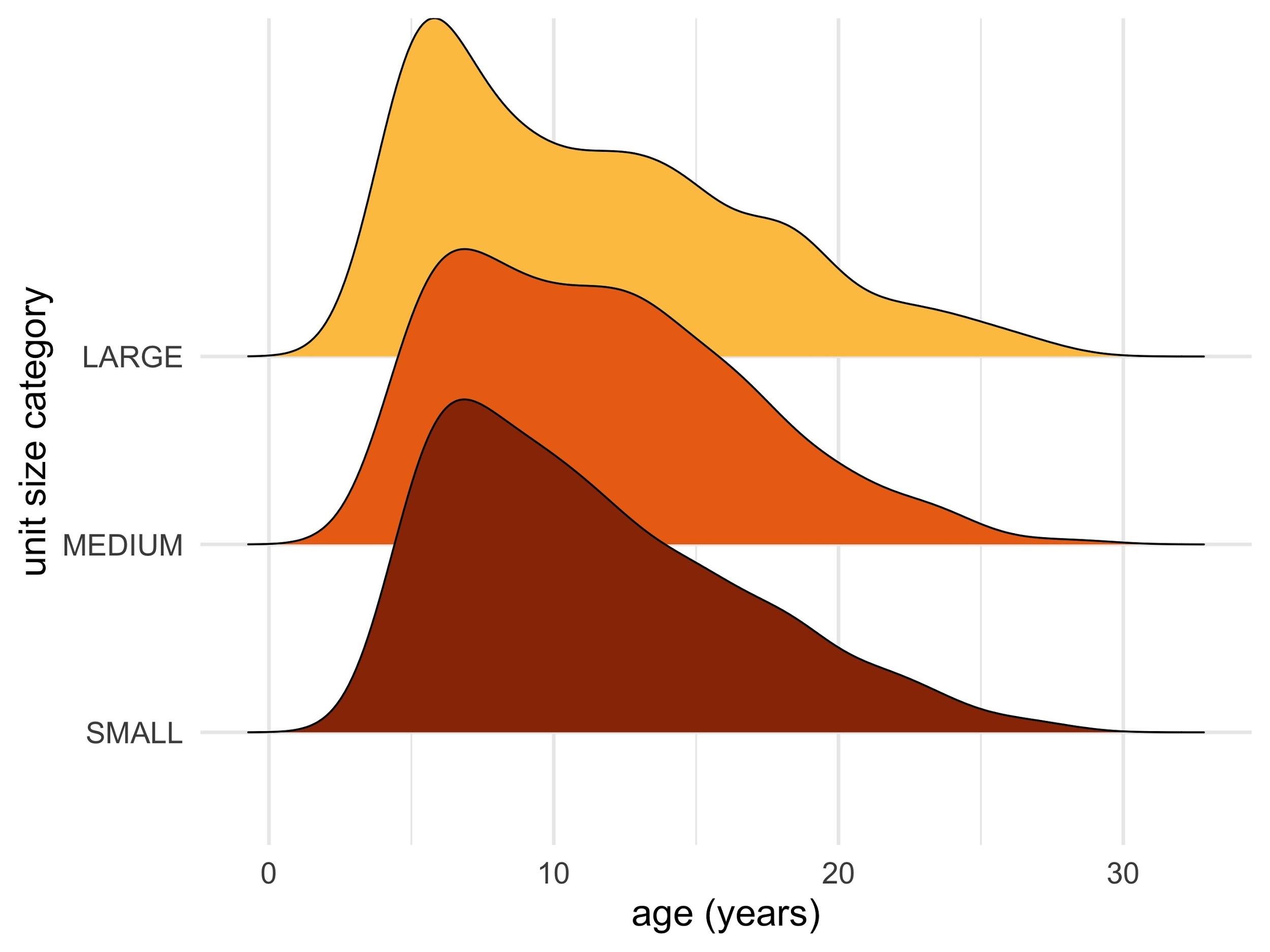

**Figure S2. Larger units are skewed towards having younger females.** The number of female-years represented in the dataset across female ages. Small unit females are labeled in red, mid-sized in orange, and large in yellow.

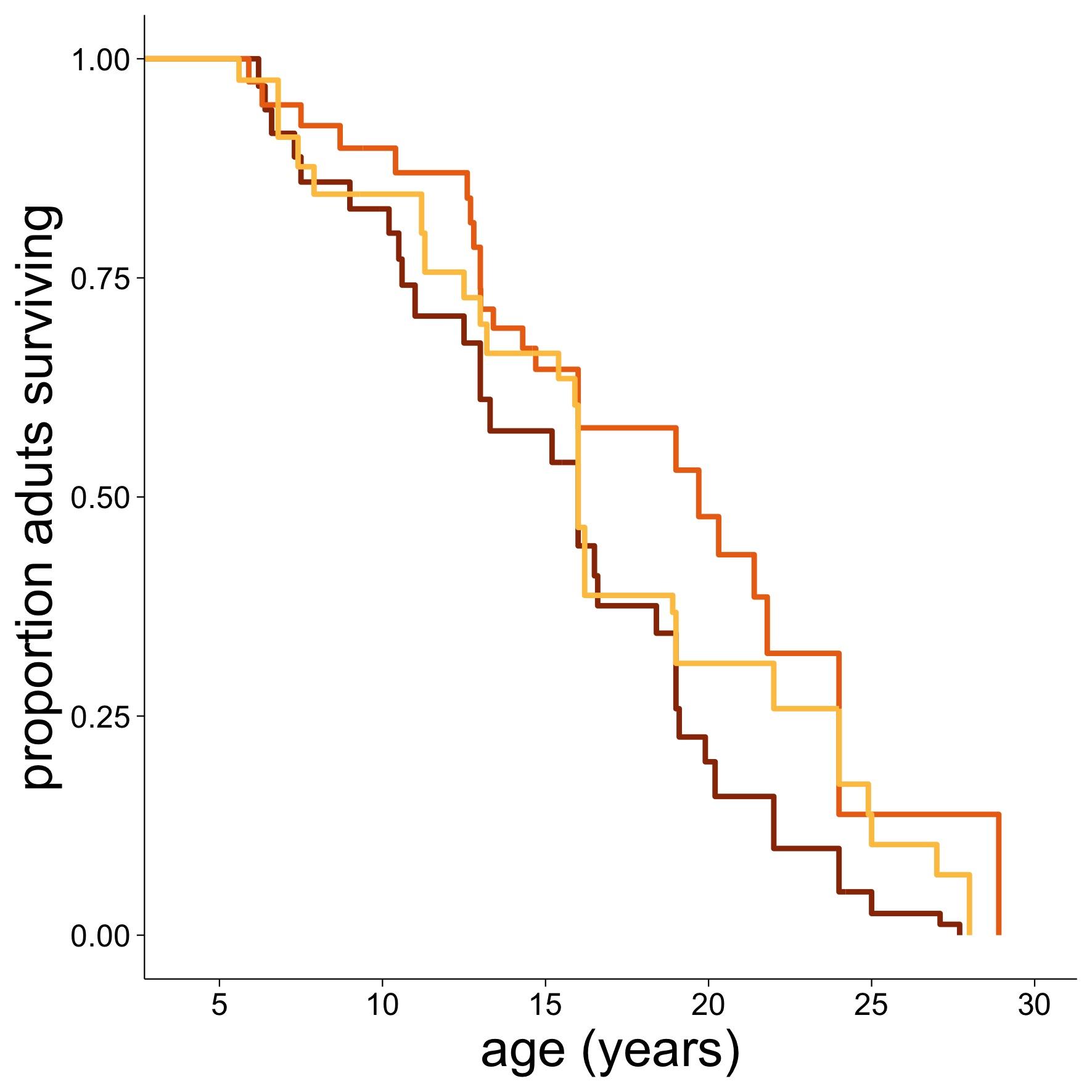

**Figure S3. Females in mid-sized units have the highest survival.** A survival curve of adult females at each unit size category. Small unit females are labeled in red, mid-sized in orange, and large in yellow. Note that unit size is a time-dependent variable: females can switch between unit size categories throughout their lifetimes as unit size changes.

**Table S1.** Results from a generalized linear mixed model examining the influence of unit size on mortality in adult female geladas.

| **Outcome** | **Predictor** | **Estimate** | **Std. Error** | **z-value** | **P-value** |
| --- | --- | --- | --- | --- | --- |
| Female mortality | (Intercept) | -2.97 | 0.32 | -9.16 | 2.0 x 10^-16^ |
|  | # females | -9.39 | 4.78 | -1.97 | 0.049 |
|  | # females^2^ | 6.78 | 3.27 | 1.59 | 0.112 |
|  | Female age | 30.07 | 7.06 | 4.26 | 2.1 x 10^-5^ |
|  | Female age^2^ | 4.12 | 4.06 | 1.01 | 0.311 |
|  | # males | -0.03 | 0.15 | -0.22 | 0.824 |
|  | **Random Effects** | **Std. Dev.** | **N** |  |  |
|  | Individual | 0.62 | 200 |  |  |
|  | Unit | 0.00 | 38 |  |  |
|  | Year | 0.10 | 15 |  |  |
| Reproductive performance | (Intercept) | -4.13 | 0.13 | -32.42 | 2.0 x 10^-16^ |
|  | # females | -17.96 | 8.02 | -2.24 | 0.025 |
|  | # females^2^ | -18.14 | 7.69 | -2.36 | 0.018 |
|  | Female age | -24.35 | 7.62 | -3.19 | 0.001 |
|  | Female age^2^ | -7.40 | 8.29 | -0.89 | 0.372 |
|  | # males | 0.20 | 0.06 | 3.26 | 0.001 |
|  | **Random Effects** | **Std. Dev.** | **N** |  |  |
|  | Individual | 0.00 | 188 |  |  |
|  | Unit | 0.00 | 40 |  |  |
|  | Year | 0.00 | 13 |  |  |

**Table S2.** Results from two linear mixed models predicting the duration of interbirth intervals (IBIs). The first model includes IBIs following all surviving offspring, while the second includes only IBIs that were uninterrupted by takeovers.

| **Outcome** | **Predictor** | **Estimate** | **Std. Error** | **z-value** | **P-value** |
| --- | --- | --- | --- | --- | --- |
| IBIs (including takeovers, n=187) | (Intercept) | 941.4 | 25.8 | 36.49 | 2.0 x 10^-16^ |
|  | # females | 492.6 | 242.6 | 2.03 | 0.044 |
|  | # females^2^ | 388.9 | 226.4 | 1.72 | 0.088 |
|  | Female age | -499.4 | 237.1 | -2.11 | 0.037 |
|  | Female age^2^ | 242.5 | 222.3 | 1.09 | 0.277 |
|  | Infant sex (M) | 68.8 | 31.54 | 2.18 | 0.031 |
|  | **Random Effects** | **Std. Dev.** | **N** |  |  |
|  | Individual | 131.6 | 104 |  |  |
|  | Unit | 0.00 | 28 |  |  |
| IBIs (no takeovers, n=82) | (Intercept) | 822.7 | 34.9 | 23.55 | 2.0 x 10^-16^ |
|  | # females | 39.2 | 230.3 | 0.17 | 0.867 |
|  | # females^2^ | 376.8 | 224.8 | 1.68 | 0.107 |
|  | Female age | -487.9 | 203.2 | -2.40 | 0.019 |
|  | Female age^2^ | 465.6 | 191.3 | 2.43 | 0.018 |
|  | Infant sex (M) | 131.7 | 37.8 | 3.49 | 0.001 |
|  | **Random Effects** | **Std. Dev.** | **N** |  |  |
|  | Individual | 162.7 | 66 |  |  |
|  | Unit | 64.4 | 22 |  |  |

**Table S3.** Results from a mixed-effects Cox proportional hazards model of infant survival.

| **Outcome** | **Predictor** | **Estimate** | **Std. Error** | **z-value** | **P-value** |
| --- | --- | --- | --- | --- | --- |
| Infant mortality | # females | 0.35 | 2.11 | 0.17 | 0.87 |
|  | # females^2^ | 6.09 | 1.91 | 3.20 | 0.001 |
|  | # males | 0.06 | 0.13 | 0.42 | 0.667 |
|  | Infant sex (M) | 0.34 | 0.22 | 1.54 | 0.120 |
|  | **Random Effects** | **Std. Dev** | **N** |  |  |
|  | Mom | 0.02 | 160 |  |  |
|  | Unit | 0.30 | 32 |  |  |
|  | Year | 0.02 | 13 |  |  |

**Table S4.** Results from Poisson generalized linear mixed model predicting the number of unit takeovers.

| **Outcome** | **Predictor** | **Estimate** | **Std. Error** | **z-value** | **P-value** |
| --- | --- | --- | --- | --- | --- |
| # takeovers | (Intercept) | -2.23 | 0.22 | -10.00 | 2.0 x 10^-16^ |
|  | # females | 7.50 | 2.47 | 3.04 | 0.003 |
|  | # females^2^ | 5.38 | 2.24 | 2.41 | 0.016 |
|  | # males | -0.13 | 0.13 | -1.02 | 0.308 |
|  | **Random Effects** | **Std. Dev.** | N |  |  |
|  | Unit | 0.00 | 46 |  |  |
|  | Year | 0.00 | 14 |  |  |

**Table S5.** Results from three generalized linear mixed models predicting infant mortality across its potential causes.

| **Outcome** | **Predictor** | **Estimate** | **Std. Error** | **z-value** | **P-value** |
| --- | --- | --- | --- | --- | --- |
| Infanticide | (Intercept) | -2.96 | 0.41 | -7.27 | 3.7 x 10^-13^ |
|  | # females | 3.48 | 4.28 | 0.81 | 0.416 |
|  | # females^2^ | 10.81 | 3.72 | 2.91 | 0.004 |
|  | **Random Effects** | **Std. Dev** | **N** |  |  |
|  | Unit | 0.88 | 31 |  |  |
| Maternal death | (Intercept) | -2.60 | 0.21 | -12.60 | 2.0 x 10^-16^ |
|  | # females | -2.48 | 3.49 | -0.71 | 0.477 |
|  | # females^2^ | 5.25 | 3.38 | 1.55 | 0.120 |
|  | **Random Effects** | **Std. Dev** | **N** |  |  |
|  | Unit | 0 | 31 |  |  |
| Unknown | (Intercept) | -2.39 | 0.19 | -12.82 | 2.0 x 10^-1^ |
|  | # females | 2.34 | 3.72 | 0.63 | 0.529 |
|  | # females^2^ | -1.07 | 3.71 | -0.29 | 0.775 |
|  | **Random Effects** | **Std. Dev** | **N** |  |  |
|  | Unit | 0 | 31 |  |  |

**Table S6.** Model averaged results from 1000 mixed effects Cox proportional hazards models estimating the influence of unit size on female death. For each model, ages were calculated by drawing a random date between the minimum and maximum dates of birth for each female.

| **Outcome** | **Predictor** | **Estimate** | **Std. Error** | **z-value** | **P-value** |
| --- | --- | --- | --- | --- | --- |
| Adult female mortality | # females | -8.02 | 3.93 | 2.04 | 0.041 |
|  | # females^2^ | 6.70 | 3.70 | 1.81 | 0.070 |
|  | # males | -0.07 | 0.14 | 0.48 | 0.630 |

**Table S7.** Results from a generalized linear mixed model examining the influence of unit size on reproductive performance. Here, reproductive performance was modeled within 6-month bins.

| **Outcome** | **Predictor** | **Estimate** | **Std. Error** | **z-value** | **P-value** |
| --- | --- | --- | --- | --- | --- |
| Reproductive performance | (Intercept) | -8.02 | 0.14 | -59.36 | 2.0 x 10^-16^ |
|  | # females | -8.81 | 3.39 | -2.60 | 0.009 |
|  | # females^2^ | -8.84 | 3.49 | -2.53 | 0.011 |
|  | Female age | -12.29 | 3.53 | -3.48 | 0.0005 |
|  | Female age^2^ | -1.56 | 3.66 | -0.43 | 0.670 |
|  | # males | 0.17 | 0.07 | 2.43 | 0.015 |
|  | **Random Effects** | **Std. Dev.** | **N** |  |  |
|  | Individual | 0.00 | 188 |  |  |
|  | Unit | 0.00 | 34 |  |  |
|  | Success Bin | 0.49 | 26 |  |  |
